# Non-invasive spinal electro-magnetic stimulation (SEMS): a tool for evaluation and modulation of lower limb spinal-muscular transmission in healthy adults

**DOI:** 10.1101/382341

**Authors:** Victor L. Arvanian, Hayk Petrosyan, Chuancai Zou, Cynthia Leone, Mohammad Zaidi, Wei Hou, Asrat Tesfa, Magda Fahmy, Mark Kaufman, Sue A. Sisto

**Affiliations:** Northport Veterans Affairs Medical Center, 79 Middleville Road, Bld. 62, Northport, NY 11768.; Stony Brook University, Department of Neurobiology and Behavior, Life Sciences Building, Stony Brook, NY 11794.; Department of Physical Therapy, School of Health Technology and Management, Stony Brook University, Stony Brook, NY 11794.; Department of Family, Preventative and Medicine, Stony Brook University, Stony Brook, NY, 11794; Department of Rehabilitation Science, School of Public Health and Health Professions, University of Buffalo, Buffalo, NY, 14214.

## Abstract

**Objective:** Our earlier electrophysiological recordings using animal models revealed diminished transmission through spared fibers to motoneurons and leg muscles after incomplete spinal cord injury (SCI). Administration of spinal electro-magnetic stimulation (SEMS) at specific parameters induced transient improvement of transmission at neuro-muscular circuitry in SCI animals. In the current human study, we sought translate this knowledge to establish optimal parameters of SEMS for (i) neurophysiological evaluation via Compound Motor Action Potential (CMAP); and (ii) modulation at neuro-muscular circuitry via H-reflex and M-wave response in 12 healthy adults.

**Methods:** SEMS application was with a coil positioned over T12-S1 spinal levels. SEMS-evoked CMAP-responses were wirelessly measured simultaneously from biceps femoris (BF), semitendinosus (ST), vastus lateralis (VL), soleus (SOL), medial gastrocnemius (MG) and lateral gastrocnemius (LG) muscles. We also examined effects of SEMS trains on H-reflex and M-wave responses. H-reflexes and M-waves were measured simultaneously from SOL, MG and LG muscles and evoked by peripheral electrical stimulation of tibial nerves before and after each SEMS session.

**Results:** Spinal levels for SEMS application to evoke CMAP-responses in corresponding muscles and amplitude/latency of these responses have been established. SEMS applied over L4-S1 spinal levels at 0.2 Hz rate for 30 min induced facilitation of H-reflexes and M-responses. Facilitation lasted for at least 1 hour after stopping SEMS and was associated with a decrease in threshold intensity and leftward shift of recruitment curve for H-reflex and M-wave. SEMS did not alter TMS-evoked responses in hand muscles.

**Conclusion:** SEMS is a novel, non-invasive approach for sustained neuromodulation of H-reflex and M-wave responses in triceps surae muscle group. The parameters of SEMS application established in this study for evaluation and neuromodulation of neural pathways innervating leg muscles in healthy individuals may be used as a reference for neurophysiological evaluation and long-lasting plasticity of the lower limb spino-neuromuscular circuitry in individuals with SCI.

## Introduction

Our recent intracellular recordings from individual axons [1] and motoneurons [2] revealed declined excitability in spared axons, impaired propagation of action potentials though these axons and thus diminished transmission to motoneurons and then to hindlimb muscles following chronic SCI in adult rats. Administration of spinal electro-magnetic stimulation (SEMS) trains induced strengthening of transmission at spino-neuromuscular circuitry in chronic SCI rats [3, 4]. The current study was designed to establish parameters of SEMS for neurophysiological evaluation and plasticity at neuro-muscular circuitry innervating various leg muscles in humans. In this initial phase, we have examined effects of SEMS in healthy adult individuals to obtain reference measures for further and on-going studies in SCI individuals.

In human physiology the electromyography (EMG), the soleus Hoffmann (H)-reflex and muscle (M) response have been traditionally used for monitoring the extent of muscle activity and neurophysiological diagnostics [5–7]. Another important neurophysiological diagnostic parameter is nerve-to-muscle conduction, which has been previously tested by either percutaneous [8], or direct needle stimulation [9] of spinal nerve roots. The remarkable ability of the electro-magnetic coil to deliver stimuli noninvasively and painlessly through skin and tissue to the deeply located peripheral nerves, has prompted several investigators to use this approach for physiological testing of the nerve-to-muscle circuitry [10–12]. This same coil has more recently been applied at thoracic spinal levels (SEMS) to evoke compound muscle action potentials (CMAPs) in leg muscles [13–15] and conditioning of these responses with peripheral electrical stimulations [14] has been examined.

Electro-magnetic stimulation is based on the principle of electromagnetic induction of an electric field through intact tissue to underlying structures systems [16–17]. Transcranial electro-magnetic stimulation (TMS), since its initial introduction about three decades ago [18], has been widely used for diagnostic applications and repetitive TMS was found to alter excitability at cortico-motor circuitry [17, 19–20]. Responses evoked by TMS, however, are reliably recorded from arm but not leg muscles, particularly in human participants with neurological impairments, including spinal cord injury [21]. Consistent with these reported results of human studies, our recent experiments using animal models revealed that TMS-evoked responses could be reliably recorded from the hind limb muscles only in the naïve animals, but not in adult rats with SCI [22]. However, responses evoked by electro-magnetic stimulation at spinal thoracic and lumbar levels could be reliably recorded from hind limb muscles in rats with chronic SCI [1]. In fact, EMG recordings from the hind limb muscles, conducted in parallel with intracellular electrophysiological recordings from individual motoneurons and axons in lumbar spinal segments, revealed that spinal electromagnetic stimulation is an excellent non-invasive approach for evaluation and modulation of transmission in spinal and spino-neuromuscular circuitry in adult rats [1, 4, 22].

Various types of functional outcomes induced by repetitive SEMS applied at different spinal levels, using different parameters and frequencies, have been reported as well. High-frequency stimulation using the EMS coil positioned over T11-T12 vertebrae induced involuntary bilateral locomotor-like movements in healthy individuals [13]. Lumbar repetitive magnetic stimulation was reported to reduce spastic tone of the lower limbs in spinal cord injury and multiple sclerosis patients [23, 24]. SEMS has been used as a treatment for urinary frequency and urge incontinence [25] as well as to suppress detrusor contraction [26]. SEMS has been used effectively to stimulate the spinal nerves in spinal cord injury individuals, resulting in some improvement of several vital functions [15, 27–29]. Research has also demonstrated greater effects of magnetic stimulation versus electrical stimulation in inhibition of detrusor hyperactivity [30]. Another study reported that repetitive magnetic stimulation at spinal levels, in combination with motor training, induced an acute and persistent decrease of low back pain and was more beneficial that motor training alone [31].

However, the CMAP responses recorded from various leg muscles and evoked by SEMS applied at different spinal levels have not been systematically examined and compared. Effects of SEMS on H-reflex and M-responses of the leg muscles have been understudied.

The first objective of the current study was to examine the ability of single-pulse SEMS to deliver electrical excitation to a variety of leg muscles in healthy adults. Using a wireless EMG system, the SEMS-evoked responses were recorded simultaneously from biceps femoris (BF), semitendinosus (ST), vastus lateralis (VL), and soleus (SOL), medial gastrocnemius (MG) and lateral gastrocnemius (LG) muscles in both legs. The second objective was to examine the ability of SEMS applied for 30 mins. to affect spino-neuromuscular plasticity: we compared M-wave and H-reflex responses that were recorded simultaneously from triceps surae muscles (SOL, MG and LG), before, immediately after the termination of SEMS, and 1 hour afterwards. We have also examined whether SEMS applied repetitively would alter TMS responses recorded from hand muscles. Some of these findings have been reported in abstract form [32].

## Material and methods

### Participants

Twelve healthy adult volunteers with no known history of neurological disorders, four male veterans (mean age 57.7 ± 7.5 years) participated in the study at the Northport VA Medical Center and 4 females (mean age 33.75 ± 9.67 years) and 4 males (mean age 26 ± 2.16 yeas) participated in the study at the Stony Brook University. All participants were screened for inclusion/exclusion criteria, gave written informed consent to participate in this study and completed up to 3 sessions. Participants were excluded from the study if they had any metal implants, any implanted electrical devices, any medications that could raise seizure threshold, cardiac conditions, history of syncope or concussion with loss of consciousness, or ringing in the ears. All females were provided a pregnancy screen where negative results allowed participation. All participants were advised not to exercise on the day of the study. This study was approved by the local Institutional Review Board (IRB) at Stony Brook University and Northport VA Medical Center and was conducted in accordance with the Declaration of Helsinki.

### Overview of Study Design

Blood pressure and heart rate were measured before and after administration of each SEMS session. First, TMS induced responses in the hand muscles were measured followed by baseline measurements of M-wave and H-reflexes recruitment curves of the leg muscles using peripheral nerve stimulation of the tibial nerve (see below). After the baseline (i.e. prior SEMS application) measurements were taken, a 30 minute session of SEMS applied in repetitive trains, was administered (see below). After the SEMS administration, M-wave and H-reflex recruitment curves and TMS induced responses were again collected for comparison with the pre-SEMS baseline measurements. To evaluate the possible long lasting effects of SEMS, the same measurements were retaken 1 hour post-SEMS administration in a subset of participants (n=4). At the Northport VAMC the study design did not include TMS administration. This sequence was repeated once a week for 3 weeks to total 3 visits.

### Electrophysiological recordings

Electrophysiological recordings were performed using Delsys Trigno wireless EMG system (Delsys Inc. Natick, MA). After standard skin preparation, wireless Trigno surface sensors were placed over the muscle belly near the motor point and aligned with the muscle striations. Lower limb recording electrodes were placed at the beginning of each session and were secured in place for the entire session, without removal. During the first session, the optimal position for placement of the recording electrodes was determined for each participant and was used for the follow-up sessions. For repeatability of EMG electrode placement for subsequent days, measurements were taken from the knee crease with a tape measure to the EMG electrode location and supported further with photographs of the electrode placement. For peripheral nerve stimulation followed by SEMS, recording electrodes were placed on the soleus (SOL), medial gastrocnemius (MG), lateral gastrocnemius (LG), vastus lateralis (VL), semitendinosis (ST) and biceps femoris (BF) muscles bilaterally. We recorded from all these leg muscles during SEMS but only triceps surae (SOL, MG and LG) for evaluation of H- and M-responses. For the TMS testing, recording electrodes were placed on first dorsal interosseous (FDI) and abductor digiti minimi (ADM) muscles. The EMG signals were recorded via Digidata 1440 digitizer (Molecular Devices, AXON CNS) at 5 kHz sampling rate using Clampex 10.6 software and analyzed off-line using Clampfit 10.6 software. The EMG responses were averaged and peak to peak amplitude from the mean response was calculated for analysis.

### Transcranial magnetic stimulation

Participants were comfortably seated in a chair with target arms well supported and completely relaxed. MagStim 200^2^ magnetic stimulator (Jali Medical, Inc. Woburn, MA) with D70^2^ magnetic coil attached to a coil stand was used for stimulation (see Fig 1). TMS was delivered to the optimal scalp position to elicit maximal responses in contralateral first dorsal interosseous (FDI) and abductor digiti minimi (ADM) muscles. After determining resting threshold intensity required to evoke responses of at least 50 μV in 3 out of 5 stimulations the stimulator output was set 20% above threshold intensity and at least 10 responses were recorded for analysis. TMS evoked responses were recorded before SEMS and after for each participant at Stony Brook University (n=8).

**Figure 1.**
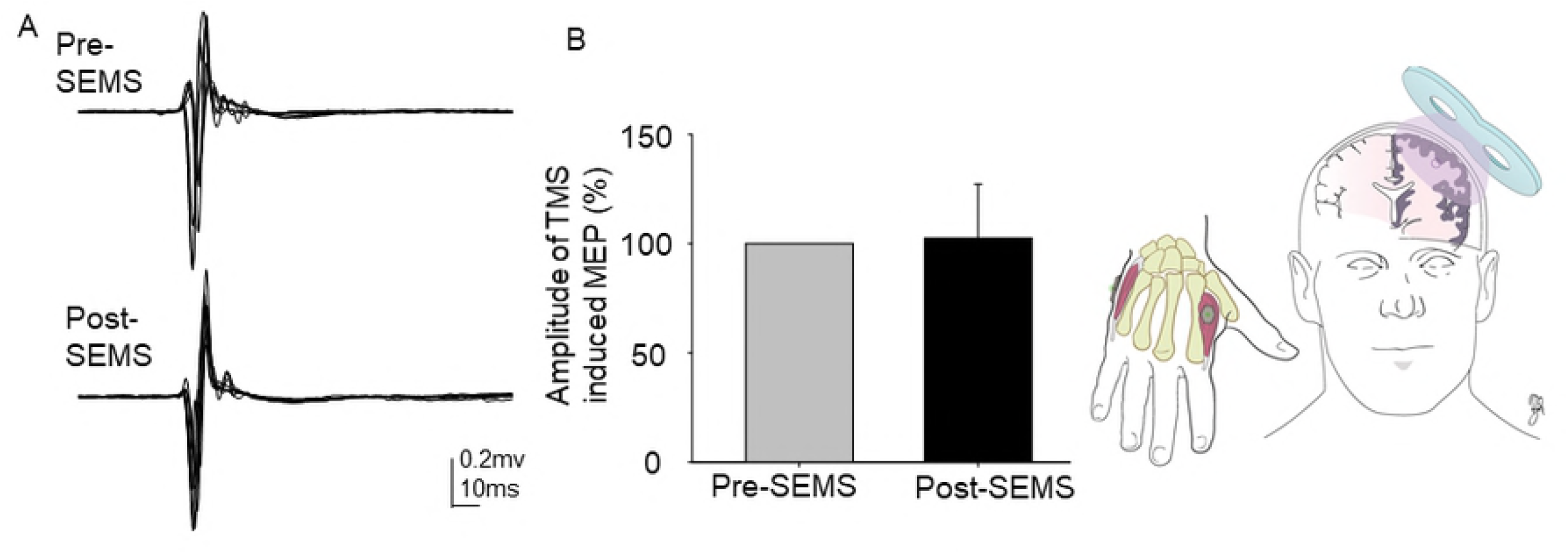
Photos are of sample participant during the experiments demonstrating the position of recording electrodes. (A) receiving SEMS administration over spinal levels and peripheral electrical stimulation over tibial nerves of both legs. (B) receiving TMS administration over the left motor cortex while recording from right hand muscles.

### Peripheral nerve stimulation, H-reflex and M-wave recordings

For peripheral nerve stimulation participants were placed in the prone position (Fig 1). All participants were instructed to refrain from voluntary contraction of the leg muscles and there was no background contraction during recordings. The M-wave and H-reflex responses were measured before and after SEMS application in all 12 participants. Peripheral electrical stimulation was delivered using Digitimer DS7A (Digitimer North America, LLC) constant current stimulator with bipolar gold-plated bar electrode (Natus Neurology, Inc.) placed over tibial nerve at the popliteal fossa. The optimal position for the stimulating electrode was based on the H-reflex and M-responses elicited and the popliteal stimulating electrode was secured in place for the duration of the session. The H-reflex and M-waves were recorded in all three triceps surae muscles using 1ms duration square-wave pulse delivered with 0.2 Hz frequency. Beginning with a low intensity stimulus that was then gradually increased until the threshold intensity for H-reflex was determined. The stimulus intensity continued to be increased in small increments (1mA) until the H maximum responses and later the M maximum responses were acquired. These data allowed for the plotting of the full stimulus/response recruitment curve. At each stimulus intensity, at least 6-10 responses were recorded. For analyses of the effects of SEMS on the recruitment curve the responses recorded from the soleus muscle were used. Six of the triceps surae EMG responses were averaged to measure the peak-to-peak amplitude for both the M-wave and H-reflex at each intensity. The data were tabulated for the baseline pre- and post-SEMS and for both left and right legs and used in the statistical analysis.

### Spinal electromagnetic stimulation (SEMS)

Participants were placed in the prone position with comfortable support of a pillow under the abdomen when preferred. SEMS was administered using MagStim 200^2^ magnetic stimulator (Jali Medical, Inc. Woburn, MA) with figure-of-8 D70^2^ magnetic coil delivering monophasic pulses at 0.2Hz frequency. The coil was always positioned with the handle towards the head to standardize the direction of electro-magnetic stimulation. The optimal spinal level and stimulus intensity were determined in their first session. To accomplish this, a total of 6-10 single pulses were delivered to each spinal level from T12 to S1 and the stimulus intensity with 10% increments ranged from 40% to 80% was applied until EMG recordings from the greatest number of muscles were visible. After the optimal spinal level and SEMS intensity were determined, they were used to deliver a 30 minutes of single pulse SEMS that was applied repetitively with 0.2 Hz frequency for each of the three sessions. Spinal levels used for repetitive SEMS ranged from L4-L5 with 60-80% of stimulus intensity. Six of all muscle consecutive responses have been collected and averaged at each intensity and spinal level, then the peak-to-peak amplitude were measured and tabulated for statistical analysis. The latency was measured from the beginning of stimulation to the onset of the response.

### Statistical Analysis

SigmaPlot 11.0 software (Systat Software, San Jose, CA) and SAS v9.3 (the SAS Institute, Cary, NC) were used for statistical analyses. For each participant the maximum H-reflex and M-response amplitudes before SEMS were normalized to 100% and the percent change for each response was calculated after SEMS. Similarly, the threshold intensity for H-reflex before SEMS was considered as 1 and used to normalize applied intensities before and after SEMS. To compare the measured characteristics of H-reflex and M-wave for pre-SEMS vs post-SEMS, paired t-test or Wilcoxon Signed Rank test were used and the results were considered statistically significant for p < 0.05. Data are presented as means ± standard errors.

To analyze the recruitment curves of H-reflex and M-wave, the ascending part of the stimulus/response recruitment curve for SOL muscle has been fitted using 3-parameter sigmoid function (SigmaPlot 11.0) with following equation for each individual [33]. Parameters such as the slope of the recruitment curve and stimulus intensity needed to obtain 50% of the maximum responses (S50) were obtained by analyses of the computer-fitted recruitment curves.

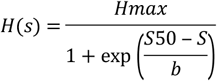

Where S is the stimulus intensity, Hmax is maximum amplitude of H-reflex. S50 is stimulus intensity needed to obtain 50% of Hmax, and b is the steepness of the curve. In addition, we calculated the slope of the curve at S50 using following equation

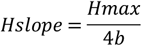

The M-wave recruitment curves were analyzed using the same equations. The amplitudes were calculated as peak to peak amplitude and intensities were normalized to pre-SEMS threshold intensity for H-reflex. i.e. for each participant/curve pre-SEMS threshold intensity for H-reflex was considered as motor threshold (MT). Because of substantial difference in actual amplitude of H- and M-responses among subjects the amplitudes were normalized to 100% of H-Max and M-Max, respectively, for each subject. All parameters analyses were grouped from each subject first and then averaged among subjects. Statistical analyses of the fitted curves were performed using SAS ver. 9.3 (the SAS Institute, Cary, NC). Descriptive statistics (e.g. means and standard deviations) of all above mentioned parameters were calculated for pre-, post- and pre-post SEMS change. The average SOL H-reflex and M-wave recruitment curves were generated using average curve parameters ± standard errors (e.g. Max, b, and S50). Three-way repeated measure ANOVA models (rmANOVA) were used to determine the effects of SEMS (pre-vs post-SEMS). Since the effects of SEMS was the primary study aim, SEMS (pre-vs post-SEMS) remained in the final models regardless of its statistical significance. The p values were then calculated based on the F tests of rmANOVA for all parameters. Due to the small sample size and the exploratory nature of this study, the primary purpose of the analysis was to examine mean trends. The conclusions will not be purely based on a significant p values [34]. Adjustment for p values depends on the number of tests and will increase the type II error [35]. Therefore, the p values were not adjusted for multiple tests [36] and any p values less than 0.05 were considered statistically significant. The correlations between M-wave and H-reflex parameters were estimated and tested for pre-, post and pre-post change of SEMS.

## Results

### SEMS-evoked CMAP responses

In this study we have examined the non-invasive electromagnetic stimulation (SEMS) administered at spinal levels in healthy participants. SEMS was administered at different spinal levels (T12-S1) and at different stimulus intensities (40 % - 80%). Fig 2 includes representative responses recorded from biceps femoris (BF), vastus lateralis (VL), semitendinosus (ST), medial gastrocnemius (MG), lateral gastrocnemius (LG) and soleus (SOL) muscles from one participant. Not surprising, the administration of SEMS at different spinal levels results in the activation of muscles differently.

**Figure 2.**
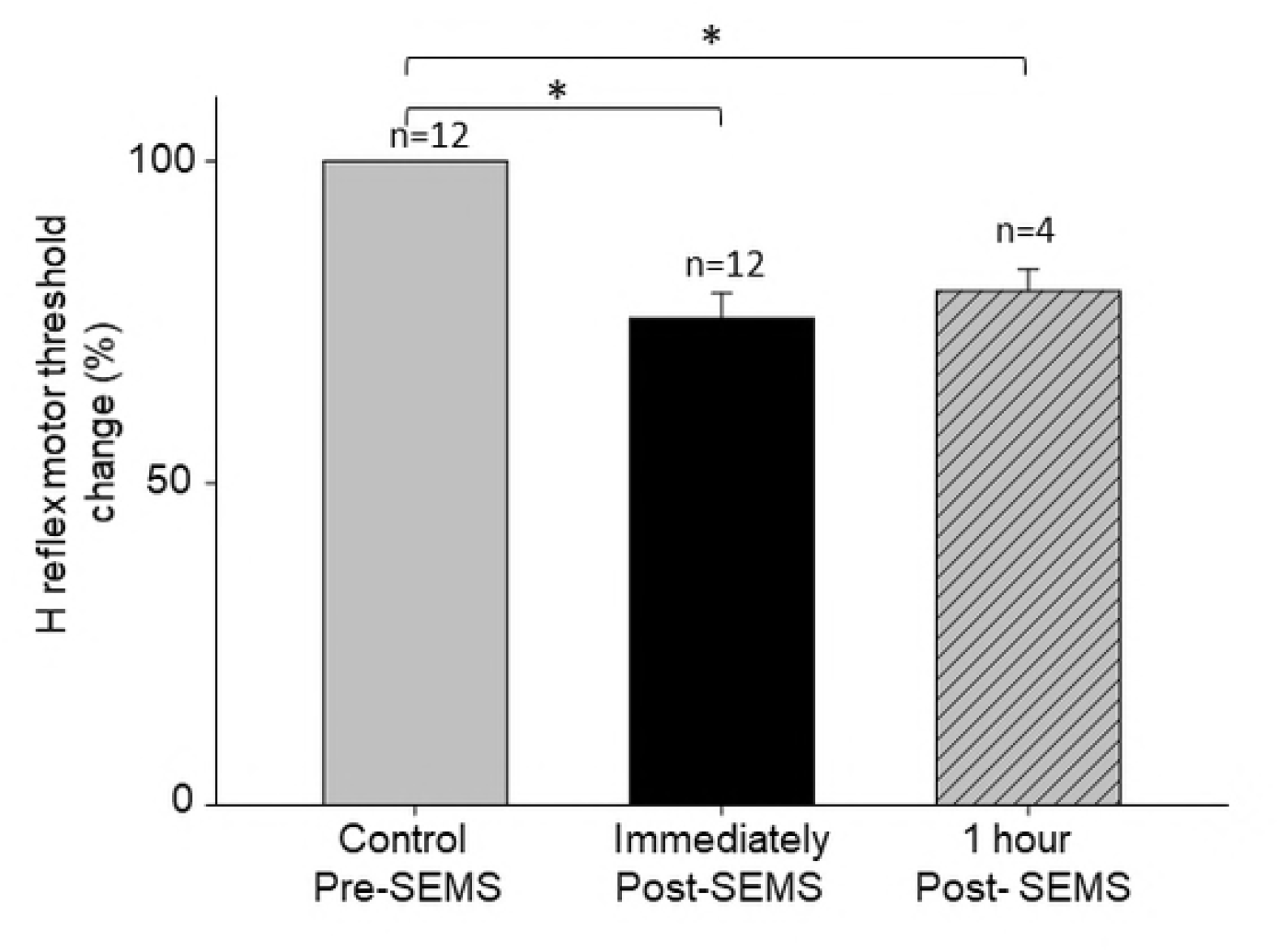
Spinal electromagnetic stimulation (SEMS) was able to evoke compound muscle action potential (CMAP) responses in several leg muscles in healthy participants. Representative responses recorded from vastus lateralis (VL), biceps femoris (BF), semitendinosus (ST), soleus (SOL), lateral gastrocnemius (LG) and medial gastrocnemius (MG) muscles are presented. Note different muscles exhibit different size and latencies in response to SEMS. Diagram illustrating the position of the SEMS stimulation coil and recording electrodes.

SEMS administration at T12-L1 levels evoked responses predominantly in BF and VL muscles (Fig 2). When SEMS was administered at low lumbar levels (L3-L5), it resulted in activation of all triceps surae muscles (MG, LG and SOL), as well as BF and ST muscles in some participants (Fig 2). Amplitude and latencies of these responses were different depending on the spinal level of stimulation and the recorded muscle. These results indicate that SEMS administration at specific spinal level results in activation of spinal different pathways to the muscles.

With the same stimulation intensity, the optimal spinal level to evoke CMAP responses in the triceps surae muscle group (MG, LG, SOL) was the L5-S1 spinal levels. The mean amplitude for the SOL muscle at 70% of the maximum coil intensity was 0.15 ± 0.02 mV whereas the MG and LG muscles showed 0.14 ± 0.3 mV and 0.10 ± 0.02 mV amplitude respectively (n=7). For the VL muscle, the optimal spinal level was T12-L1. The mean amplitude for the VL muscle at 70% of maximum coil intensity was 0.14 ± 0.07 mV. For the BF and ST muscles, the optimal spinal stimulation level was L1-L3. The mean amplitude for BF muscle at 70% of maximum coil intensity was 0.03 ± 0.01 mV and ST was 0.05 ± 0.01 mV respectively.

Latencies of CMAP responses were also different depending on the muscle. Latency for the VL muscle responses evoked from L1 SEMS was 11.8 ± 1.1 ms, 13.5 ± 1.3 ms for BF muscle and 12.8 ± 1.9 ms for the ST muscle. Latency of responses evoked by coil positioned over L5 for triceps surae muscles showed slightly longer latencies, i.e. 15.3 ± 0.53 ms for MG; 14.7 ± 0.45 ms for LG and 15.4 ± 0.54 ms for SOL. The CMAP responses from the same muscles exhibited different latencies and different amplitudes when the SEMS coil was moved caudally. In the same participant, the CMAP responses recorded from SOL muscle showed decreased latency and increased amplitude as the SEMS coil was adjusted caudally. For example, in one participant the latency at L1 was 21.2 ms but at the L5 level was 18.1 ms and the amplitude was 0.01mV at L1 and 0.21mV at L5 respectively.

### Effects of single pulse SEMS with repetitive trains on M-wave and H-reflex amplitudes

We have examined effect of SEMS administration for 30 min on the M-wave and H-reflex responses. We measured the M-wave and H-reflex response in all triceps surae muscles before and after administration of SEMS train (see methods). The H-reflex and M-wave responses elicited by electrical stimulation of the tibial nerve at the popliteal fossa can be observed in Fig 3A. The responses recorded simultaneously from the SOL, MG and LG muscles, varied among participants. Fig 3B shows the M-wave and H-reflex responses recorded from the same participant immediately after SEMS administration using same current intensity as in Fig 3A. Both H- and M-responses exhibited facilitation in all three muscles after SEMS. Similar facilitation after SEMS administration was observed in all 12 participants tested. Since the SOL muscle has been examined extensively in the literature, the motor threshold (MT), or the minimum stimulus intensity that produced a minimal measurable H-response (about 30 μV in at least 5 of 10 trials) in the SOL muscle was used for further analyses. This effect of SEMS on H-reflex and M-response was associated with changes in threshold intensities for both H- and M-responses (see below). There was no change in the latencies of both H-and M responses after SEMS administration. Before SEMS the mean latency for SOL muscle of M response was 8.15 ± 0.32ms and of H was 28.5 ± 0.42ms; after SEMS administration latency for M was 8.18 ± 0.32ms and for H was 28.8 ± 0.41ms (p > 0.05; n=12).

**Figure 3.**
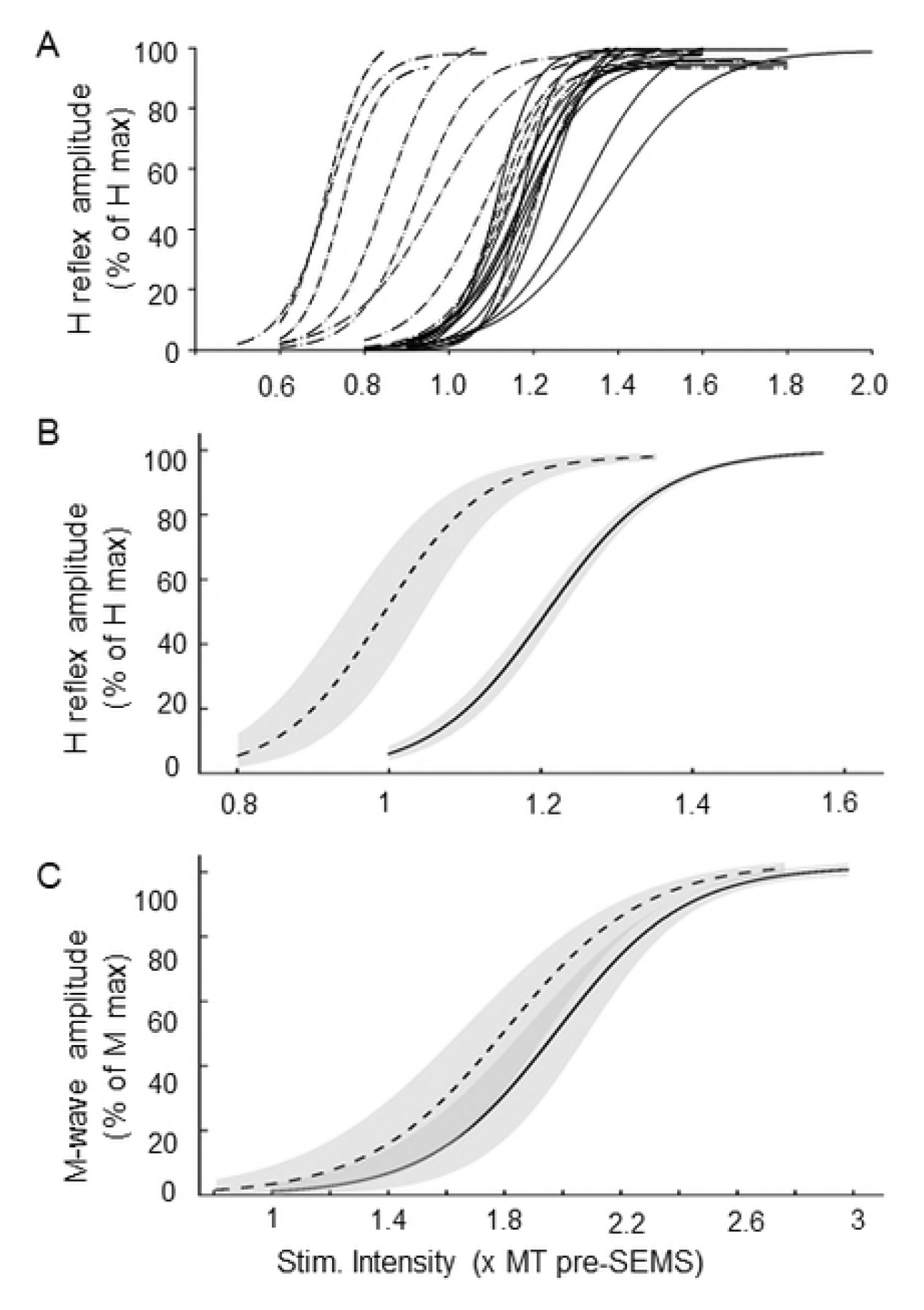
Modulation of H-reflex and M-wave by SEMS train in healthy participants. After repetitive SEMS (0.2 Hz for 30 min) amplitude of both H-reflex and M-wave responses show marked facilitation in all three soleus (SOL), Medial Gastrocnemius (MG) and Lateral Gastrocnemius (LG) muscles. (A) Representative traces of H-reflex and M-wave responses recorded from SOL, MG and LG muscles before (A) and after SEMS train (B) from same participant evoked with same current intensities stimulating tibial nerve (approx. 20% above motor threshold). Diagram illustrating the position of the stimulation electrode for tibial nerve stimulation and recording electrodes for H-reflex and M-wave.

In two participants, we examined whether changes in positioning of the figure of-eight shaped electro-magnetic coil over human vertebrae would affect amplitude of SEMS-evoked CMAP responses, or SEMS-induced modulation of M- and H-responses. Rotating the coil 180°, with the handle positioned towards the legs instead of towards the head, while keeping its center at same spinal level, did not induce changes in the amplitude of SEMS-evoked responses. Additionally, it did not affect the ability of SEMS administration to modulate the M-wave and H-reflex responses. These results are consistent with previous reports indicating that (i) current flowing in a clockwise or anticlockwise directions did not induce a difference in the amplitude of the responses in humans [14], (ii) rotation of the coil 180°, did not induce significant changes in the effects of SEMS on synaptic transmission in animal models [1], (iii) neither it affected movement of the surgical rods [37]. However, all data represented are from participants receiving SEMS with the coil cable positioned cephalad.

### Effects of single pulse SEMS administration with repetitive trains on Motor Threshold (MT) of M-wave and H-reflex

We have also systematically examined the effects of SEMS administration on the threshold intensity required to evoke H-reflex and M-wave responses. Fig 4A demonstrates representative traces of M-wave and H-reflex responses, recorded at different stimulus intensities in SOL muscle from one participant before and after administration of SEMS. After SEMS administration (for 30 min at 0.2 Hz), the threshold intensity for both H- and M-responses was significantly lower. Full intensity/response recruitment curve for the SOL muscle for the same participant is presented in Fig 4B (participant 1). After SEMS administration, there was a leftward shift (to lower intensity) for both responses. A similar shift of the recruitment curve and decrease of threshold intensity was observed in all participants. Representative recruitment curves from randomly chosen two other participants (participant 2 and participant 3), are presented in Fig 4B as well. Results revealed that administration of SEMS for 30 minutes induced a substantial leftward shift of the recruitment curve for both H- and M-responses; however, this shift was different among participants (Fig 4). SEMS-induced changes of H-reflex and M-wave responses in SOL muscle were compared in each participant first and then grouped among participants. The mean threshold intensity for H-reflex was 12.4 ± 1.1 mA pre-SEMS vs 8.8 ± 0.7 mA post-SEMS, i.e. 73 ± 3% of pre-SEMS control (P < 0.05, n = 7). The mean threshold intensity for M-wave was 17.4 ± 1.6 mA pre-SEMS vs 12.1 ± 1.2 mA post-SEMS, i.e. 70.84 ± 4.31% of pre-SEMS control (P < 0.05, n = 7).

**Figure 4.**
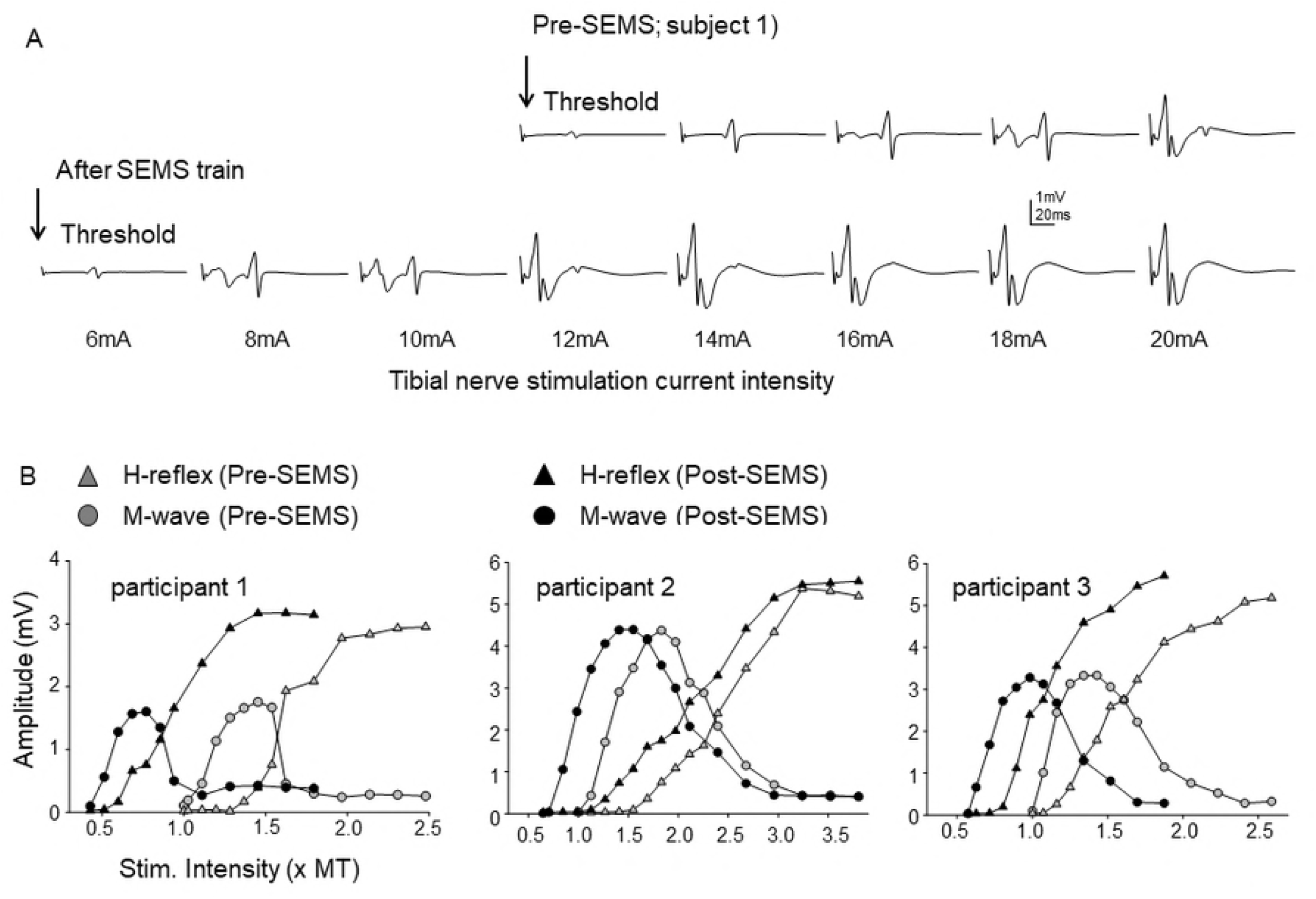
Effect of SEMS train on recruitment curve of H-reflex and M-wave responses. (A) Representative traces of H-reflex and M-wave responses recorded from SOL muscle in one participant (participant 1) before and after SEMS. After SEMS there is a dramatic change in the current required to evoke H-reflex. Representative traces of H-reflex and M-wave are presented for each stimulation current intensity before and after SEMS. (B) Representative stimulus/response recruitment curves for H-reflex and M-wave for the same (participant-1) and two other (participants 2 and 3) participants before and after SEMS demonstrating a leftward shift of both H-reflex and M-wave curves.

### Statistical analyses of the recruitment curves for H- and M-responses

A summary of the results demonstrating changes in characteristics of M-wave and H-reflex responses in SOL muscle following administration of SEMS train (30 min, 0.2 Hz) is presented in Fig 5. In order to generalize results of the SEMS effects on the H- and M-recruitment curves in all participants it was important to account for the differences in MT intensity and amplitude of H- and M-responses among participants. Therefore, the stimuli intensities were normalized to the pre-SEMS values of the MT; and amplitudes of H- and M-responses were normalized to 100% of maximum responses pre-SEMS, respectively, for each participant. The rising phase of recruitment curves for H-reflex and M-wave was fitted using sigmoid function and analyzed (see methods).

**Figure 5.**
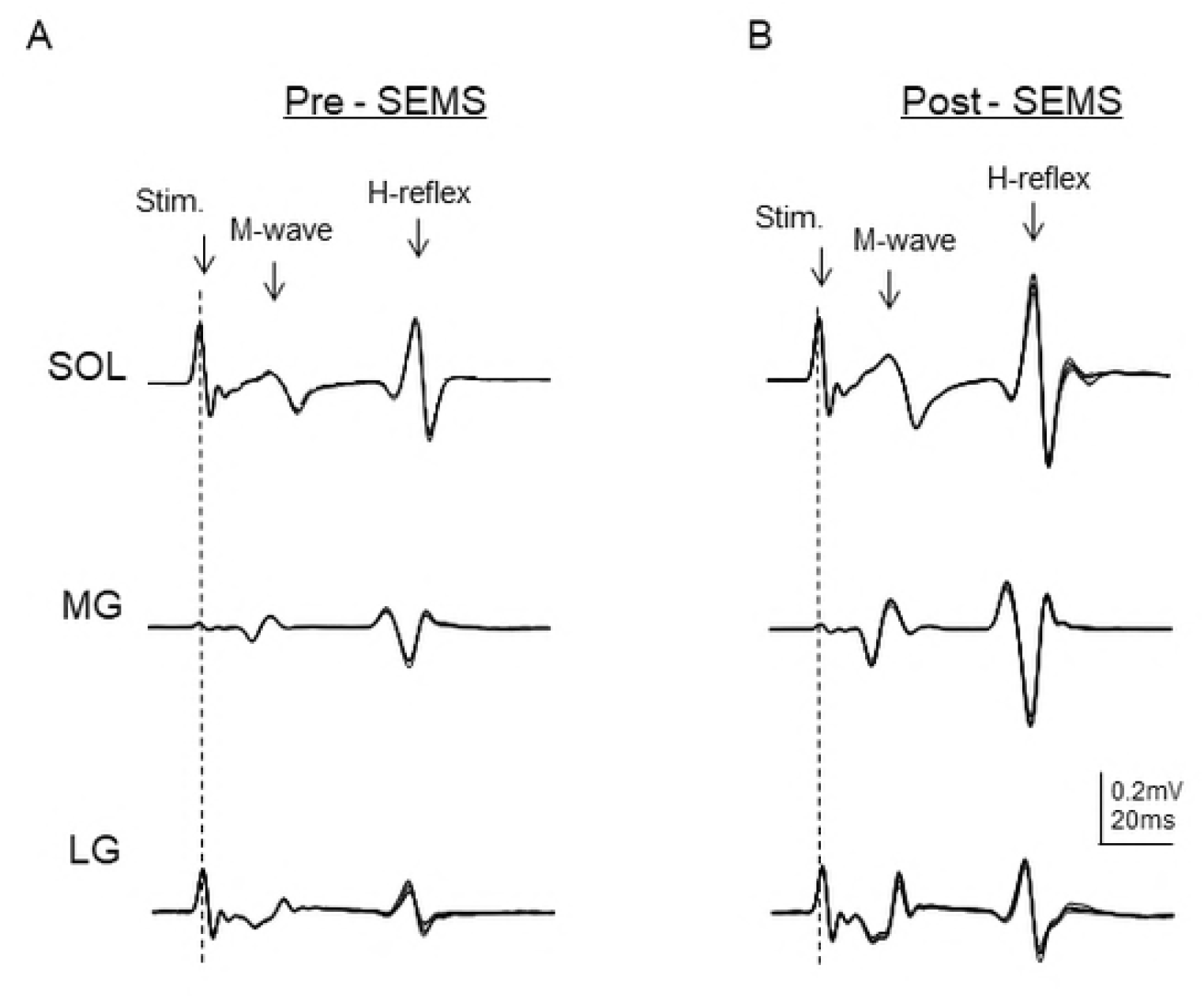
Computer-generated analyses to summarize results of the effects of SEMS trains on recruitment curves for M-wave and H-reflex responses among all participants. (A) Plot demonstrating H-reflex recruitment curves using three parameter sigmoid fits for each participant. Solid lines demonstrate sigmoid fits before SEMS and doted lines demonstrate H-reflex sigmoid fits after SEMS for each participant. (B, C) Averaged curves for H-reflex and M-wave before (solid lines) and after SEMS administration (dotted lines) respectively. Shaded area represents standard error.

Analyses of the parameters of fitted M-wave curve are summarized in Table 1. The slope at 50% M-max was flatter after SEMS compared to before SEMS (157.51±76.42 pre-SEMS vs 139.78±53.95 post-SEMS, p=0.03, Mean ± SD). S50 parameter before SEMS was significantly greater than S50 after SEMS (1.98±0.44 pre-SEMS vs 1.81±0.59 post-SEMS, p=0.03, Mean ± SD). However, M-max, b, and intensity range were not significantly different before compared to after SEMS.

**Table 1.** Descriptive Statistics of Parameters for M Wave, Pre and Post SEMS

| Parameter | Pre<br>N=19 | Post<br>N=19 | Change<br>(Post-Pre SEMS)<br>N=19 | p-value<sup>1</sup> |
| --- | --- | --- | --- | --- |
|  | Mean±SD<br>Median (range) | Mean±SD<br>Median (range) | Mean±SD<br>Median (range) |  |
| **M-max** | 101.8±10.0<br>100.09 (93.81, 140.82) | 103.03±8.5<br>100.64 (94.17, 132.95) | 1.3±4.6<br>0.73 (-7.87, 11.22) | 0.45 |
| **b (Steepness)** | 0.2±0.2<br>0.2 (0.07, 0.8) | 0.2±0.2<br>0.16 (0.12, 0.87) | 0.02±0.04<br>0.01 (-0.06, 0.08) | 0.10 |
| **Mslope (Slope at 50% M-max)** | 157.5±76.4<br>142.5 (44.21, 330.42) | 139.8±54<br>159.03 (40.37, 207.57) | -17.7±55.5<br>-4.07 (-143.94, 59.91) | 0.03 |
| **Intensity corresponding to M-max (x MT)** | 3.0±0.6<br>2.8 (2.42, 4.14) | 2.8±0.78<br>2.6 (1.79, 4.43) | -0.2±0.3<br>-0.2 (-1.03, 0.40) | 0.05 |
| **Intensity range (Max - Min) (x MT)** | 2.0±0.6<br>1.8 (1.42, 3.14) | 2.0±0.7<br>1.7 (1.16, 3.5) | -0.04±0.2<br>-0.05 (-0.45, 0.36) | 0.60 |
| **S<sub>50</sub> (Intensity corresponding to 50% M-max) (x MT)** | 2.0±0.4<br>1.9 (1.51, 3.36) | 1.8±0.6<br>1.7 (0.94, 3.35) | -0.2±0.3<br>-0.1 (-0.75, 0.22) | 0.03 |

<sup>1</sup>Testing the significance of pre-post change based on F-tests of rmANOVA.

Analyses of the parameters of fitted H-reflex curve are summarized in Table 2. The stimulus intensity corresponding to H-max before SEMS was higher compared to after SEMS (1.57±0.14 pre-SEMS vs 1.35±0.32 post-SEMS, p=0.02, Mean±SD). The S50 parameter before SEMS was also significantly higher than after SEMS (1.21±0.06 pre-SEMS vs 0.99±0.21 post-SEMS, p=0.02, Mean±SD). However, H-max, b, Slope at 50% H-max and Intensity Range are not significantly different before and after SEMS.

**Table 2.** Descriptive Statistics of Parameters for H Reflex by Pre and Post SEMS

| Parameter | Pre<br>N=18 | Post<br>N=18 | Change (Post-Pre)<br>N=19 | P-value<sup>1</sup> |
| --- | --- | --- | --- | --- |
|  | Mean±SD<br>Median (range) | Mean±SD<br>Median (range) | Mean±SD<br>Median (range) |  |
| **H-max** | 99.9±4.2<br>98.7 (94.52,107.55) | 98.4±3.5<br>98.2 (92.46,104.60) | -1.5±3.7<br>-0.8 (-10.23, 5.60) | 0.26 |
| **b (Steepness)** | 0.1±0.02<br>0.1 (0.04, 0.11) | 0.1±0.02<br>0.1 (0.04,0.11) | -0.01±0.02<br>-0.01 (-0.05,0.02) | 0.24 |
| **Hslope (Slope at 50% H-max)** | 352.7±105.1<br>338.6(216.07, 616.72) | 397.2±108.3<br>381.4(221.45, 582.10) | 44.5±108.2<br>36.05 (-167.28,305.21) | 0.29 |
| **Intensity corresponding to H-max (x MT)** | 1.6±0.1<br>1.56 (1.40,1.83) | 1.4±0.32<br>1.5 (0.77, 1.78) | -0.2±0.3<br>-0.2 (-0.72, 0.18) | 0.02 |
| **Intensity range (Max - Min) (x MT)** | 0.6±0.14<br>0.56 (0.40, 0.83) | 0.6±0.2<br>0.6 (0.21, 0.90) | -0.02±0.1<br>-0.02(-0.35, 0.26) | 0.30 |
| **S<sub>50</sub> (Intensity corresponding to 50% H-max) (x MT)** | 1.2±0.06<br>1.2 (1.11, 1.37) | 1.0±0.2<br>1.1 (0.54, 1.29) | -0.2±0.2<br>-0.1 (-0.62, 0.10) | 0.02 |

<sup>1</sup>Testing the significance of pre-post change based on F-tests of rmANOVA.

In order to understand whether SEMS induced changes in H-reflex and in M-wave are comparable, we have also analyzed correlation between SEMS-induced changes in the parameters of H-reflex and M-wave recruitment curves. As shown in Table 3, the SEMS-induced changes in the intensity corresponding to M-max and H-max, as well as S50 parameters were significantly correlated for M-wave and H-reflex responses (both r>0.07, p <0.001). However, important to note that Hslope and Mslope at 50% of max and steepness of the curves were not correlated (r=0.09, p=0.71 and r=0.08, p=0.74 respectively).

**Table 3.** Correlations between parameter estimates of H-reflex and M-wave

| Parameter | Correlation Coefficient r<br>(p-value) |  |  |
| --- | --- | --- | --- |
|  | Pre | Post | Change |
| **Normalized M-Max and H-Max** | -0.09<br>(0.72) | -0.12<br>(0.62) | 0.46<br>(0.06) |
| **B (Steepness)** | -0.05<br>(0.84) | 0.25<br>(0.32) | 0.08<br>(0.74) |
| **Hslope and Mslope (at 50% of Hmax and Mmax)** | -0.07<br>(0.7810) | 0.27<br>(0.2838) | -0.09<br>(0.71) |
| **Intensity corresponding to M-max and H-max (x MT)** | -0.24<br>(0.3338) | 0.31<br>(0.2047) | 0.75<br>(0.0004) |
| **Intensity range (Max - Min) (x MT)** | -0.24<br>(0.3338) | 0.13<br>(0.6198) | 0.41<br>(0.09) |
| **S<sub>50</sub> (Intensity corresponding to 50%) (x MT)** | -0.17<br>(0.50) | 0.47<br>(0.047) | 0.70<br>(0.001) |

### Modulation of H-reflex by SEMS train is long-lasting and sustained after stop of SEMS

We have also examined whether the effect of SEMS are long-lasting, i.e. how long the observed threshold changes were sustained by measuring H-reflex before, immediately after SEMS and after 1 hour post stopping of SEMS (Fig 6). The threshold intensity required for H reflex was still significantly lower after 1 hour post SEMS administration;79.8 ± 3.3 % post-SEMS compared to 100% pre-SEMS (n=4; p<0.05). Importantly, the decrease in the MT intensity following SEMS application, however, was not associated with the changes in Hmax. Hmax amplitude was 1.96 ± 0.19 mV before and 1.91 ± 0.12 mV after SEMS application (n = 8, p = 0.156).

**Figure 6.**
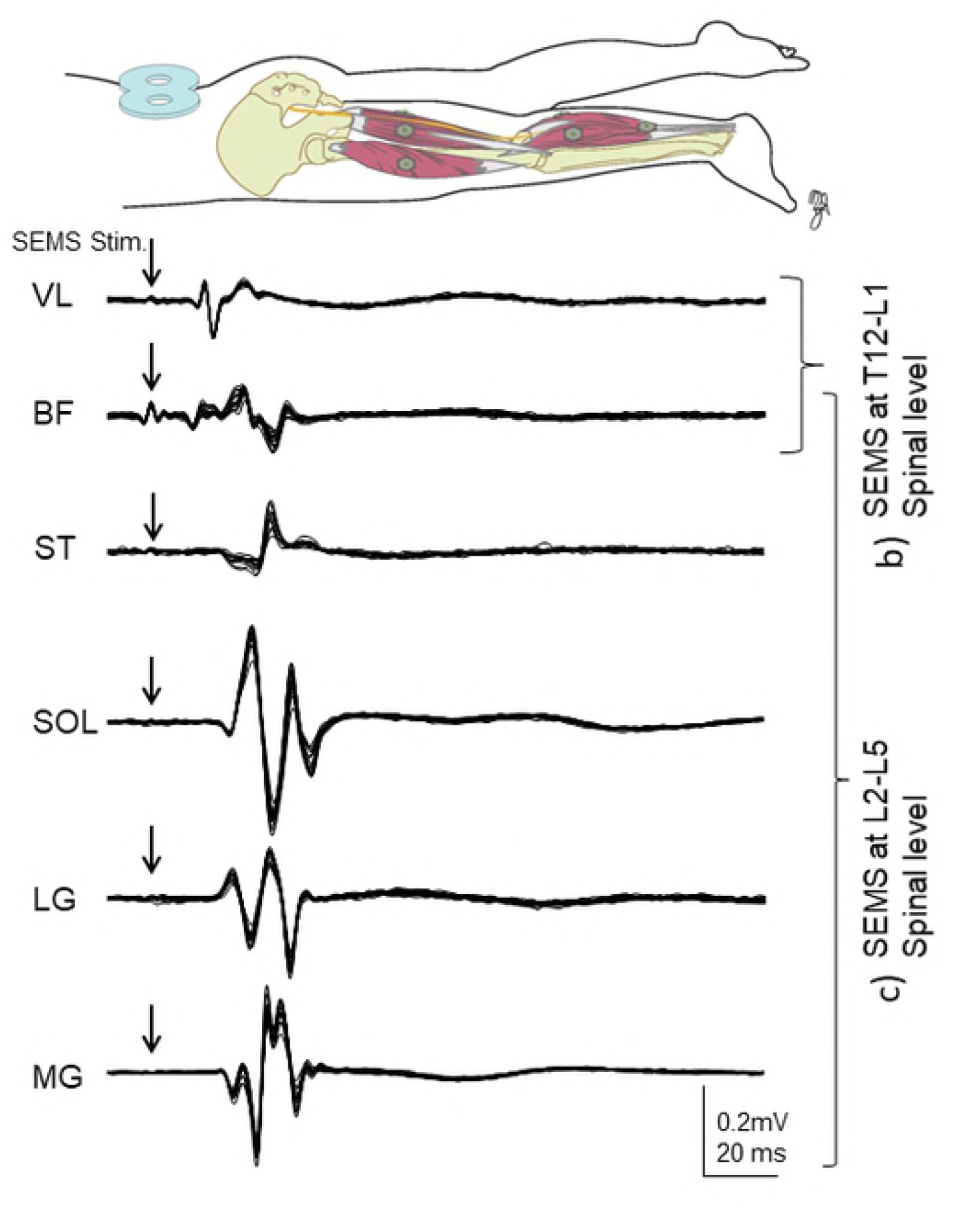
Long lasting effect of SEMS on the H-reflex. Summary of results demonstrating change in the stimulation current intensity required to elicit the minimal H-reflex after SEMS administration for all participants. Summary of results are presented as percent change of control for each participant, where control (before SEMS) represent 100%. Effect of SEMS in decreasing threshold intensity lasts for at least 1 hour after stop of SEMS.

### Effects of SEMS on TMS evoked responses

We have also examined whether SEMS administration may affect TMS induced responses. We recorded TMS evoked responses from hand FDI muscle before and after SEMS administration. Representative traces recorded from FDI muscle before and after SEMS in the same participant presented in Fig 7A. We did not observe any changes in TMS evoked responses after SEMS. Percentage change of TMS evoked MEP responses is presented in Fig 7B. There was no significant change in the amplitude of MEP recorded from FDI muscles after SEMS i.e. 102.6 ± 24.1 % (n=8; p > 0.05). These results suggest that an observed effect of SEMS on H-reflex and M wave responses are local and do not involve supraspinal systems.

**Figure 7.**
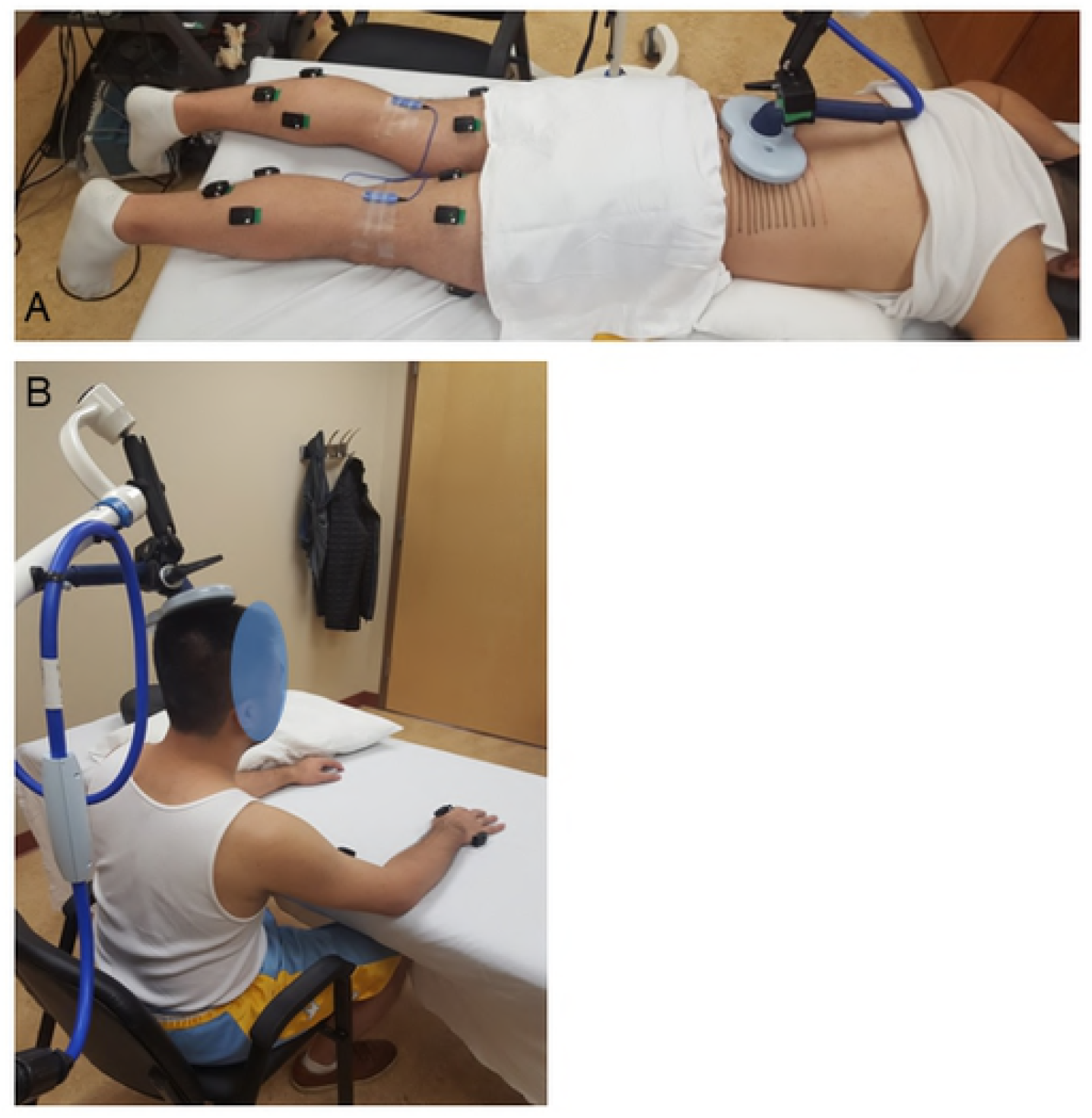
Repeated SEMS administration does not affect the TMS-evoked responses recorded from hand muscles. (A) Representative TMS evoked motor evoked potentials (MEP) recorded from first dorsal interosseous (FDI) muscle from one participant before and after SEMS administration. (B) Summary of results demonstrating no change in the amplitude of MEPs recorded from FDI muscles in response to TMS after SEMS administration. Results are presented as percent change of control for each participant. Diagram illustrating the position of TMS stimulation coil and recording electrodes.

## Discussion

There are two major outcomes from this study. (1) Parameters for SEMS stimulation and characteristics of CMAP responses recorded simultaneously from several leg muscles have been established; results demonstrate that CMAP responses measured from different muscles in legs and evoked by single-pulse SEMS applied at different spinal levels can serve as an feasible approach for the neurophysiological evaluation of neural pathways innervating leg muscles. (2) SEMS applied as a train of repetitive pulses may modulate the amplitude of both the direct (M-wave) and reflex (H-reflex) responses recorded from SOL, MG and LG leg muscles. No previous study has compared effects of repetitive SEMS on M- and H-responses of these three sections of the triceps surae muscle group. Together these results suggest that SEMS may become a novel non-invasive and painless tool to induce long-lasting plasticity of the lower limb spino-neuromuscular circuitry.

### SEMS applied at different spinal levels is an effective non-invasive stimulation protocol for activation of distinct leg muscles

Electro-magnetic stimulation applied at the periphery i.e. over spinal roots, nerves or muscles has been recently utilized in clinical research as a new painless and noninvasive approach to deliver electrical excitation [13, 29]. Neurophysiological parameters obtained by measures of CMAP responses evoked by lumbosacral electro-magnetic stimulation are comparable with those obtained with electrical motor-root stimulation using surface electrodes [10, 38], or by inserting needle electrodes into the tissues around the lumbosacral motor roots [39, 40]. However, these types of electrical stimulation often associated with pain, discomfort, and sensory disturbance, in contrast to non-invasive and painless SEMS application. Systematic examination and comparison of CMAP responses recorded in different leg muscles and evoked by administration of SEMS at different spinal levels in same participant is understudied.

Several previous reports describe neurophysiological characteristics of CMAPs responses evoked by SEMS. For example, positioning of the coil over the C7 spine was optimal for hand muscle stimulation [41] and SEMS administration over thoracic spinal levels induced activation of leg muscles [13, 14]. Applied at lumbosacral level it was used for estimates of peripheral motor nerve conduction and diagnosis of lumbosacral motor radiculopathy [42].

Our current study examined and described the properties of CMAP responses recorded simultaneously from several muscles (soleus, MG, LG, BF, ST and VL) in both legs and evoked by positioning of electro-magnetic coil at different spinal levels (T12 to S1) using stimulus of various intensity (40% to 80% of coil maximum intensity). Our results demonstrate that SEMS administration at different spinal levels result in activation of distinct muscles with different CMAP properties. CMAP responses recorded from the same muscles were showing different latencies and different amplitudes depending on the spinal level where SEMS was administered. Presented results demonstrate that SEMS is an effective non-invasive tool that can activate distinct pathways and diverse leg muscles when applied at different spinal levels. Together these results indicate that SEMS is an appropriate approach for neurophysiological evaluation of conduction at neural pathways innervating leg muscles. The measures of amplitude and latency of CMAP responses recorded from various muscles in healthy individuals may serve as a reference for future studies examining effects of SEMS in SCI individuals or other neurological disorders.

### SEMS applied as repetitive trains is an effective tool to modulate H-reflex and M-response

Another important finding was that SEMS applied as repetitive train induced modulation of electrically evoked M-response and H-reflex in the SOL, MG, LG muscles. The H-reflex which is a response elicited by electrical stimulation of afferent (Ia sensory) fibers travels to the motoneurons pool of the corresponding muscle in the spinal cord with subsequent monosynaptic recruitment of alpha motor neurons and then through the efferent (motor) fibers to the muscle spindles [6, 43]. H-reflex and muscle M-response evoked in the SOL muscle by the tibial nerve electrical stimulation are recognized as standard tools for studying the integrity and plasticity at spino-muscular circuitry [44, 45]. Any change in H-reflex amplitude under a given condition reflects changes in the excitability of the afferent/motoneuron/efferent reflex pathways, while changes in M-wave amplitude reflect changes in excitability in the efferent fibers only.

In this study we have examined effect of SEMS on the amplitude of M- and H-responses recorded simultaneously from SOL, MG and LG muscles. Although all these three muscles belong to triceps surae muscle group, these muscles differ in function, composition, and innervation [45]. Comparison of the effects of SEMS on M- and H-responses of these muscles have not been described by other groups.

Our results demonstrate that application of SEMS for 30 mins at 0.2Hz frequency induced substantial facilitation of both M-response and H-reflex in all three SOL, MG and LG muscles. To better characterize the SEMS-induced modulation of M- and H-responses we have analyzed the effects of SEMS on the recruitment curves that have been obtained from SOL muscles. Results suggest that effects of repetitive SEMS associated with the significant leftward shift of the recruitment curves for both M- and H-responses. Effects of SEMS on H-reflex and M-wave were associated with a marked decrease in the threshold currents to evoke H- and M-responses, with no change in latencies of these responses. This effect of SEMS lasted for at least 1 hour after stopping of SEMS. The leftward shift of the recruitment curve for H-reflex and the decrease in MT current intensity to evoke the H-reflex after SEMS application was not associated with changes in the amplitude of Hmax (Figs. 4, 5), known as an estimate of the number of motor neurons (MNs)s that could be activated [46, 47]. Although the mechanisms underlying effects of repetitive SEMS on neural plasticity are yet to be discovered, SEMS-induced a leftward shift of the recruitment curve and a decrease of the MT for both M- and H-responses clearly indicate that prolonged (30 min) application of pulsed SEMS at low (0.2 Hz) frequency induces substantial changes in the excitability of spino-muscular circuitries.

### Comparison of the effects of SEMS with other types of stimulation to modulate H-reflex and M-response

Our results of the neuromodulation of M-wave and H-reflex responses induced by non-invasive SEMS application is similar to results reported by other types of stimulation techniques. For example similar modulation of M-wave and H-reflex responses was reported with transcutaneous direct current stimulation (tsDCS), where tsDCS paired with transcranial magnetic stimulation TMS induced substantial reduction in the motor threshold for both M- and H-responses [48]. Applied alone, tsDCS was found to modulate characteristics of M-wave and H-reflex as well, with type of modulation depending on applied current polarity, i.e. anodal or cathodal current application [49–51]. This type of neuromodulation has been attributed to changes in the efficacy of the Ia fiber-motoneuron synapse [52, 53], or excitability at spino-neuromuscular circuitry [51]. Modulation of synaptic transmission within the spinal cord may involve changes in several neural properties, including changes in membrane potential with a corresponding changes in synaptic activity [54, 55] and/or modulation of neuronal firing rate [56, 57]. Several reports suggest the primary role of modulation of axonal excitability in the effects of tsDCS [51, 58, 59]. Effect of tsDCS on axonal excitability was suggested to be attributed to depolarizing current interaction with axonal voltage-gated Na+ channels [60, 61]. Long-lasting effects of tcDCS on H-reflex has been suggested to be a result of triggering a self-sustained opening of ‘persistent’ Na+ channels [51], known to be able to keep the membrane potential steadily depolarized [62] and alter the distribution of proteins on the plasmatic membrane [63].

SEMS and established electrical stimulation techniques may, however, act differently on neural plasticity and have different effects within spino-muscular circuits. In contrast to electrical stimulation, it has been reported that SEMS activates deep conductive structures, produces strong muscle contractions and recruits proprioceptive afferents with minimal cutaneous recruitment of sensory fibers [64–66]. While there are some differences between SEMS and tcDCS induced neuromodulations we do not exclude a possibility that these cellular mechanisms suggested to describe the modulatory effects of the tcDCS may be involved in the reporting here neuroplasticity induced by non-invasive SEMS. Although the intrinsic processes underlying the effects of SEMS on M- and H-responses reported in this study require further examination, our results strongly suggest that SEMS is an effective and non-invasive approach to modulate function in spino-neuromuscular circuitry.

Results of our analyses demonstrate strong correlation of some parameters (i.e. Normalized Mmax and Hmax; Intensity corresponding to Mmax and Hmax; S50; intensity corresponding to 50% of max response), but lack of correlation of other parameters of H-reflex and M-wave recruitment curves such as steepness of the fitted recruitment curves, intensity range and changes of H-slope and M-slope at 50% of Hmax and Mmax. These observations suggest that SEMS may exert its modulatory action at the level of (i) spinal neural networks, including efficacy of synaptic transmission and/or neuron membrane potentials, or (ii) modulating properties of peripheral nerve, i.e. excitability of the afferent sensory and/or efferent motor axons, or (iii) occurs at both spinal and peripheral levels. Experiments using animal models are required to determine the mechanisms that underlay the SEMS induced neuroplasticity and these experiments are on-going.

In conclusion, reported here for the first time the ability of the SEMS to exert the long-lasting modulation of excitability at spino-neuromuscular circuitry. suggests a potential application of SEMS in clinics for number of spinal or peripheral nerve conditions. Thus modulating the strength of the motor and sensory inputs could lead to a change of the H-reflex characteristics [67]. SEMS could be also used for improving the diminished axonal conduction as is the case of many clinical neurological disorders [68], including SCI [2, 3], as it has been demonstrated using animal SCI models [1, 4, 22].

## Acknowledgements

The authors appreciate the technical assistance of Mohammed Harb.

